# Mechanistic modeling quantifies the influence of tumor growth kinetics on the response to anti-angiogenic treatment

**DOI:** 10.1101/136531

**Authors:** Thomas D. Gaddy, Alyssa D. Arnheim, Stacey D. Finley

**Affiliations:** Department of Chemical Engineering and Materials Science, University of Southern California, Los Angeles, California, United States of America; Department of Biomedical Engineering, Boston University, Boston, Massachusetts, United States of America; Department of Biomedical Engineering, University of Southern California, Los Angeles, California, United States of America

## Abstract

Tumors exploit angiogenesis, the formation of new blood vessels from pre-existing vasculature, in order to obtain nutrients required for continued growth and proliferation. Targeting factors that regulate angiogenesis, including the potent promoter vascular endothelial growth factor (VEGF), is therefore an attractive strategy for inhibiting tumor growth. Computational modeling can be used to identify tumor-specific properties that influence the response to anti-angiogenic strategies. Here, we build on our previous systems biology model of VEGF transport and kinetics in tumor-bearing mice to include a tumor compartment whose volume depends on the “angiogenic signal” produced when VEGF binds to its receptors on tumor endothelial cells. We trained and validated the model using published *in vivo* measurements of xenograft tumor volume, producing a model that accurately predicts the tumor’s response to anti-angiogenic treatment. We applied the model to investigate how tumor growth kinetics influence the response to anti-angiogenic treatment targeting VEGF. Based on multivariate regression analysis, we found that certain intrinsic kinetic parameters that characterize the growth of tumors could successfully predict response to anti-VEGF treatment, the reduction in tumor volume. Lastly, we use the trained model to predict the response to anti-VEGF therapy for tumors expressing different levels of VEGF receptors. The model predicts that certain tumors are more sensitive to treatment than others, and the response to treatment shows a nonlinear dependence on the VEGF receptor expression. Overall, this model is a useful tool for predicting how tumors will respond to anti-VEGF treatment, and it complements pre-clinical *in vivo* mouse studies.

## Introduction

Angiogenesis is the formation of new blood vessels from pre-existing vasculature and is important in both physiological and pathological conditions. Numerous promoters and inhibitors regulate angiogenesis. One key promoter of angiogenesis is the vascular endothelial growth factor-A (VEGF-A), which has been extensively studied and is a member of a family of pro-angiogenic factors that includes five ligands: VEGF-A, VEGF-B, VEGF-C, VEGF-D, and placental growth factor (PlGF). VEGF-A (or simply, VEGF) promotes angiogenesis by binding to its receptors VEGFR1 and VEGFR2 and recruiting co-receptors called neuropilins (NRP1 and NRP2). The VEGF receptors and co-receptors are expressed on many different cell types, including endothelial cells (ECs), cancer cells, neurons, and muscle fibers [1]. Together, VEGF and its receptors and co-receptors initiate the intracellular signaling necessary to promote vessel sprouting, and ultimately, the formation of fully matured and functional vessels. The new vasculature formed following VEGF signaling enables delivery of oxygen and nutrients and facilitates removal of waste products [2].

Regulating angiogenesis presents an attractive treatment strategy for diseases characterized by either insufficient or excessive vascularization. In the context of excessive vascularization seen in many types of cancer, inhibiting angiogenesis can decrease tumor growth. Anti-angiogenic treatment targeting tumor vascularization is a particular focus area within cancer research [3]. One anti-angiogenic drug is bevacizumab, a recombinant monoclonal antibody that neutralizes VEGF (an “anti-VEGF” drug). Bevacizumab is approved as a monotherapy or in combination with chemotherapy for several cancers, including metastatic colorectal cancer, non-small cell lung cancer, and metastatic cervical cancer [4]. In 2008, the drug gained accelerated approval for treatment of metastatic breast cancer (mBC) through the US Food and Drug Administration (FDA), based on evidence from pre-clinical studies and early phase clinical trials. Though initial clinical trials initially showed that bevacizumab improved progression-free survival (PFS), subsequent results revealed that bevacizumab failed to improve overall survival (OS) in a wide range of patients and that the drug elicited significant adverse side effects [5]. Consequently, the FDA revoked its approval for the use of bevacizumab for mBC in late 2011 [6].

The case of bevacizumab illustrates that although anti-angiogenic therapy can be effective, not all patients or cancer types respond to the treatment. This underscores the need for biomarkers that can help select patients who are likely to respond to anti-angiogenic treatment. Numerous studies have sought to identify biomarkers for anti-angiogenic treatment. Biomarkers can be used to determine which tumors will respond prior to any treatment being given (“predictive”), or to evaluate efficacy following treatment (“prognostic”) [7]. Biomarkers can also be used to determine optimal doses, to design combination therapies, and to identify resistance to therapies [8]. The concentration range of circulating angiogenic factors (CAFs), and VEGF in particular, is one possible predictor of the response to anti-angiogenic therapy [7]. Alternatively, expression of angiogenic receptors such as NRP1 and VEGFR1 on tumor cells, in the tumor interstitial space, or in plasma can serve as biomarker candidates [5,9]. Unfortunately, though some of these candidates are promising, a marker that predicts bevacizumab treatment outcome has not yet been validated [5,7]. In fact, relying on the concentrations of CAFs has produced inconclusive and inconsistent results [7,8,10]. Tumor growth kinetics have also been investigated as prognostic biomarkers of the response to anti-angiogenic treatment [11-15]. The most recent studies take advantage of improved imaging technology that can assess tumor volume, rather than only providing two-dimensional information [11]. The imaging analyses show that tumor growth kinetics may be a reliable indicator of treatment efficacy and are in good agreement with standardized approaches for assessing response treatment. However, utilizing tumor growth kinetics as a predictive biomarker has not been extensively studied.

Mouse models present a useful platform for cancer research, including biomarker discovery. Despite differences in the mouse and human anatomy and immune system, pre-clinical mouse studies are useful in understanding human cancer progression and response to therapy [16]. Advances in molecular biology techniques have generated relevant mouse models (i.e., patient-derived tumor models and genetically engineered models). These mouse models enable biomarker discovery for early detection of cancer [17], to identify non-responders to a particular treatment [18], and to classify tumors as being drug-sensitive or drug-resistant [19]. Excitingly, computational analyses are being combined with pre-clinical models to identify biomarkers for early detection and progression [17,19].

There is a substantial and productive history of applying computational modeling to study cancer at multiple scales, from initiation through metastasis [20-22]. The model predictions provide testable hypotheses that have been experimentally and clinically validated. Given the multiple cell types, molecular species and signaling pathways involved in angiogenesis, systems biology approaches are used to understand the dynamic ligand-receptor interactions that mediate angiogenesis and tumor growth. Systems biology studies how individual components of biological systems give rise to the function and behavior of the system and aims to predict this behavior by combining quantitative experimental techniques and computational models [23]. Our previous work and the work of others demonstrates that mathematical models complement pre-clinical and clinical angiogenesis research [8,24]. These models have been used to identify prognostic biomarkers that can predict which patients will benefit from anti-angiogenic therapies [24-26].

In this work, we use a computational systems biology model to investigate the utility of tumor growth kinetics in predicting response to anti-VEGF treatment. We make use of quantitative measurements from pre-clinical mouse studies and use those data to train the computational model. This work builds upon our previous computational model of VEGF distribution and kinetics in tumor-bearing mice [27] by changing the dynamic tumor volume to be dependent on the pro-angiogenic complexes involving VEGF-bound receptors (the “angiogenic signal”). This new element of the computational model allows us to simulate anti-VEGF treatment and predict the effect of the treatment on tumor volume. We apply the new model to identify conditions and characteristics of tumor growth that may be predictive of a favorable response to anti-angiogenic treatment. Our work contributes to the identification of validated biomarkers that could be used to determine tumors that are sensitive to anti-angiogenic treatment.

## Results

### Model construction

We have previously developed compartmental models to investigate the kinetics and transport of VEGF, a key regulator of angiogenesis [28-32]. In our previous computational model, the dynamic tumor volume was given by an exponential function and was not linked to the concentrations of pro-angiogenic species. We now address this limitation of our previous work. Specifically, we expand our previous computational model of VEGF distribution in tumor-bearing mice [28] to incorporate the effect of VEGF on tumor growth. Having the dynamic tumor volume be a function of the concentration of VEGF bound to receptors on tumor endothelial cells is a significant improvement and generates a more physiologically relevant computational tool to investigate anti-angiogenic treatment strategies.

Details regarding the model structure and molecular species are provided in the Methods Section. Here, we detail the equation for tumor growth. Tumor growth is given by an adapted Gompertz model focusing on the exponential and linear phases of the tumor, as previously described [8,33]. Thus, the differential equation for the tumor volume (termed “Tumor Growth Model 1”) is:

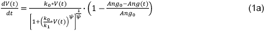

We note that equation (1a) simplifies to:

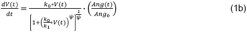

Here, *V(t)* is the tumor volume in cm^3^ at time *t, k_0_* and *k_1_* are parameters describing the rate of exponential and linear growth, respectively. The units of *k_0_* and *k_1_* are s^-1^ and cm^3^ tissue/s, respectively. The *ψ* parameter represents the transition from exponential to linear tumor growth and is unitless. The *Ang_0_* parameter represents the basal angiogenic signal (at time *t* = 0), and *Ang(t)* is the angiogenic signal at time t. The value of *Ang* at any time is calculated as the total concentration of pro-angiogenic VEGF-receptor complexes on tumor endothelial cells. This includes VEGFR1 and VEGFR2 bound to either mouse or human VEGF isoforms, with or without the NRP1 co-receptor. Thus, *Ang(t)* and *Ang_0_* have units of concentration (mol/cm^3^ tissue). The values of the tumor growth parameters (k_0_, *k_1_, ψ* and *Ang_0_*) were estimated by fitting the model to experimental data, as described in the following section.

### Model fitting

We fit the model to control data from published experimental datasets quantifying tumor volume in mice bearing MDA-MB-231 xenograft tumors without any anti-VEGF treatment [34-38]. Tumor growth is somewhat variable across the datasets, with the final tumor volume ranging from 0.8–2.5 cm^3^. The raw data used for fitting (extracted from published references; see Methods for details) are provided in the Supplementary File S1.

We used nonlinear least squares optimization to fit the model and estimate the optimal parameter values, minimizing the error between the model predictions and the experimental measurements. Specifically, we estimated the values of the tumor growth parameters, including *k_0_*, *k_1_*, *ψ*, and *Ang_0_* (see Methods section for more details). We performed the model fitting 20 times for each of the six datasets, obtaining 20 sets of optimized parameter values per dataset. Overall, the model does a good job of recreating the growth dynamics of untreated tumors (Fig 1, blue lines). One limitation is the fit to data from Volk *et al.* [38], where the model fails to capture the sigmoidal shape of the experimental tumor growth curve (Fig 1F).

**Fig 1.**
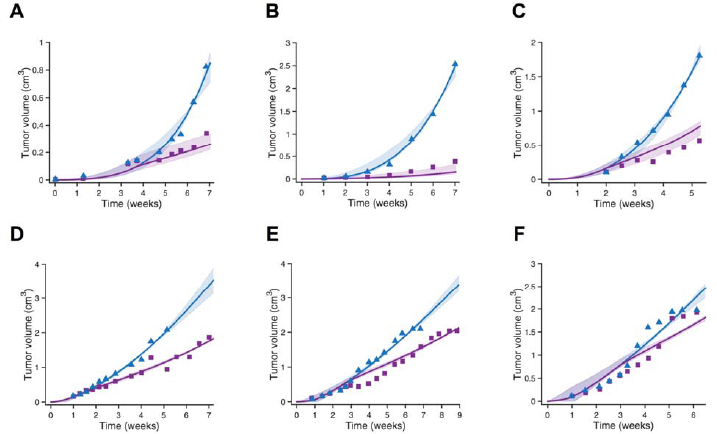
Model fit and validation using full tumor growth time course for fitting. The whole-body mouse model was used to fit measurements of tumor xenograft volumes, and the tumor growth kinetic parameters were estimated. The predicted tumor volume over time is shown for the six datasets. **A**, Roland [34]. **B**, Zibara [35]. **C**, Tan [36]. **D**, Volk [37]. **E**, Volk 2011a [38]. **F**, Volk 2011b [38]. The model is able to reproduce experimental data for tumor growth without treatment and predict validation data not used in parameter fitting. Blue triangles and purple squares are control and treatment experimental data points, respectively. Solid lines are the mean of the model results obtained from the best fits, and the shading indicates the 95% confidence interval. Note different scales on both axes.

We also explored an alternate equation for predicting tumor growth. In a separate model fitting, we augment equation (1) to include a coefficient *(C_Ang_)* that describes how dependent the tumor growth is on the concentration of the VEGF-VEGFR species (termed “Tumor Growth Model 2”):

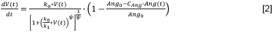

Overall, including the *C_Ang_* as another fitted parameter did not significantly improve the fit to the training data (i.e., the error is the same for both Models; Supplemental File S2). Additionally, the estimated values of *C_Ang_* spanned a fairly large range (Supplemental File S3), indicating that the value of *C_Ang_* does not significantly influence the quality of the fit to the experimental data. That is, given the available experimental data, the *C_Ang_* parameter may be non-identifiable. Therefore, we moved forward with Tumor Growth Model 1, which includes four parameters that characterize the kinetics of tumor growth (equation (1)).

### Model validation

We validated the model with data not used in the fitting. Using the same fitted kinetic parameters as the control case, we simulated the treatment regimens described in the *in vivo* mouse studies. The model does an excellent job of matching the experimental data (Fig 1, purple lines), capturing the effect of anti-VEGF treatment on tumor growth for the majority of datasets. Based on these results, the model is in agreement with experimental data of untreated tumor growth and can be appropriately validated using treatment data. Thus, our model is able to recreate the growth dynamics of untreated breast tumor xenografts in mice and can predict the tumor volume in response to anti-angiogenic treatment.

### Model fitting to early tumor growth data

We investigated whether it is possible to accurately predict the response to anti-VEGF treatment when the model fitting only includes the initial tumor growth data. We selected the datasets that included at least three tumor volume measurements prior to administration of bevacizumab (two out of the six datasets fit this criterion). We fit those initial experimental data points for the control (no anti-VEGF treatment) and validated the fitted model using the anti-VEGF treatment data. We again performed the model fitting 20 times for each dataset. The optimized model fit using only the initial tumor growth data was able to predict the tumor volume following treatment (Fig 2). Although the 95% intervals were wider in this fitting as compared to the results obtained when all of the data points were used for model fitting (see Fig 1), the newly optimized model still predicted reasonable values for the tumor size. Overall, these results demonstrate that the model can recreate both control and treatment dynamics even when parameter fitting is performed using a limited number of experimental measurements.

**Fig 2.**
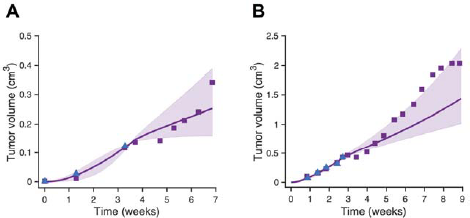
Model validation after fitting initial tumor growth data. Predicted tumor volume over time for the two datasets with at least three pre-treatment measurements for tumor volume. **A**, Roland [34]. **B**, Volk 2011a [38]. Triangles (blue) and squares (purple) are control and treatment experimental data points, respectively. Only the blue data points are used for fitting. Solid line is the mean of the predicted results obtained from the best fits, and the shading shows the 95% confidence interval on the best fits. Note different scales on both axes. Confidence intervals for the predicted tumor volumes with treatment were reasonably small and contained the experimental data points.

### Analysis of the estimated parameter values

We evaluated the optimized parameters estimated from fitting the model to all of the available control data. The estimated parameter values for the fits with the lowest errors are given in Fig 3. When fitting to the datasets from Volk et al., the model fitting and parameter estimation showed higher *ψ* values and *k_0_lk_1_* ratios than the other three datasets. Some of the estimated parameter values varied widely, even up to 3-4 orders of magnitude, while others, such as *ψ*, had a much narrower range.

**Fig 3.**
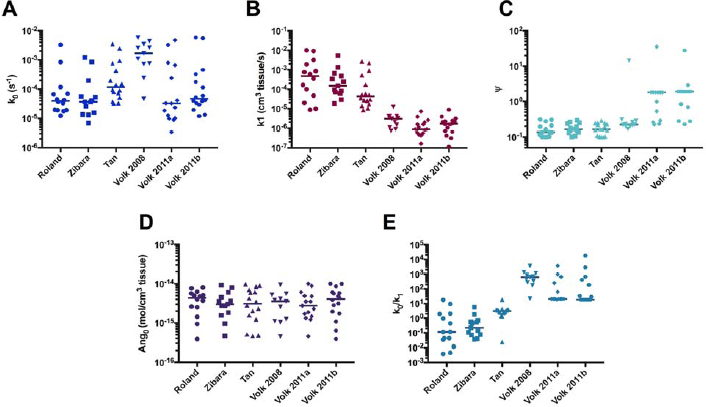
Estimated model parameters obtained from fitting. The whole-body mouse model was used to fit measurements of tumor xenograft volumes, and the tumor growth kinetic parameters were estimated. The estimated parameter values from the best fits are plotted for each dataset. **A**, *k_0_*. **B**, *k_1_*. **C**, *ψ* **D**, *Ang_0_*. **E**, *k_0_lk_1_*. Horizontal bar indicates the median of the best fits obtained from fitting the model to each dataset. Statistical comparison of the estimated parameter sets is given in Fig 5.

We performed statistical analyses to compare the estimated parameter values obtained from fitting the full set of control data to the estimated parameters obtained from fitting only the initial measurements before treatment. The analyses revealed no significant difference between the estimated parameter values obtained from the two fittings (p > 0.9 for all parameters). Thus, the fitted model obtained using only the initial time points provides reliable predictions regarding the effect of anti-VEGF treatment on tumor volume and is consistent with the optimized model obtained from fitting all of the data. Overall, the model predicts tumor growth well, even with a limited number of pre-treatment data points used for model fitting. This highlights the robustness of the model in that only three to five data points were needed in order to get a reasonable prediction for treatment response. However, for the remainder of the paper, we present simulations obtained using the optimized parameter sets estimated from fitting the full time course for the control data. Using the fitting from the full training data allows us to compare the response to treatment across all six independent experimental datasets.

### Predicting the response to anti-VEGF treatment

Having validated our model, we used the optimized parameter sets to predict the tumor volume in response to anti-VEGF treatment. We ran the model for each of the six datasets, using all 20 sets of optimized parameter values. For each set of parameters, the model was simulated for three cases: no treatment (control) and two treatment conditions (2 and 10 mg/kg bevacizumab). For the treatment cases, twice-weekly injections were simulated, starting when the tumor volume reached 0.1 cm^3^ (termed “T_start_”). We selected this volume, since it is established that this is the critical time at which tumors typically start secreting higher levels of angiogenic factors in order to recruit the vasculature necessary to support further growth (~1-2 mm in diameter). For all cases, the model was simulated for 6 weeks after *T_start_*. We used the model to predict the relative tumor volume (RTV), the ratio of the final tumor volume for the control and treatment cases:

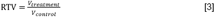

where *V_treatment_* and *V_control_* are the tumor volumes at the end of the simulation with treatment and without treatment, respectively. Thus, the RTV represents the fold-change in tumor size due to treatment, where an RTV value less than one indicates that the treatment reduced the tumor volume, compared to the control. We use the RTV value to characterize the response to anti-VEGF treatment. The predicted response to bevacizumab treatment at a dose of 10 mg/kg using the best fit parameter values is shown in Fig 4. The predicted responses for a dose of 2 mg/kg are shown in Supplemental File S4. The range of predicted RTV values indicates that certain tumors are more responsive to anti-VEGF treatment than others. In particular, the predicted RTV values obtained using fitted parameter values from fitting to data from Volk are higher than the predicted response for the other datasets. We next performed a thorough statistical comparison of the RTV and the estimated parameter values obtained in the fitting.

**Fig 4.**
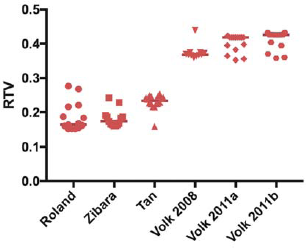
Predicted response to anti-VEGF treatment. The whole-body mouse model, including the dynamic tumor compartment whose volume is predicted using equation (1), was used to simulate bevacizumab treatment at a dose of 10 mg/kg. The relative tumor volume (RTV) predicted by the model is shown. Horizontal bar indicates the median of the predicted RTV for the best fits from each dataset.

Our statistical analysis indicates a relationship between particular kinetic parameters that characterize tumor growth and the effectiveness of treatment. We used to statistical analyses to determine whether the sets of estimated parameters or the predicted RTV values were statistically significantly different (p < 0.05) across the six datasets (Fig 5). Based on this analysis, we found that all datasets with significantly different predicted RTV values either had significantly different *k_1_, ψ*, values or *k_0_lk_1_* ratios. The *ψ* parameter represents the switch from exponential to linear growth [8], and *k_0_/k_1_* is the ratio of the exponential to linear growth rates [33]. Interestingly, there was no statistically significant difference in the estimated *Ang_0_* values, the “basal angiogenic signal”, between any of the datasets. Overall, the statistical analysis reveals that certain kinetic parameters (*k_1_, ψ*, and *k_0_lk_1_*) varied considerably between datasets and corresponded to significantly different treatment response (as indicated by the RTV value). The values of those parameters, which characterize the kinetics of tumor growth, may be used to predict the response to treatment.

**Fig 5.**
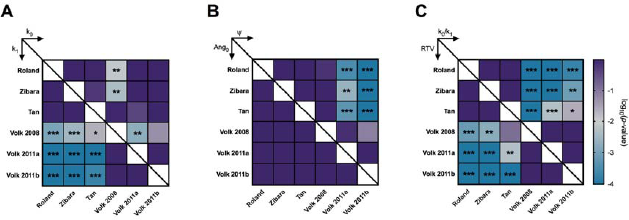
Statistical analysis of the optimized parameter sets. Standard ANOVA analysis followed by pairwise comparisons was used to determine whether the sets of optimized parameter values were statistically different. **A**, upper triangle: *k_0_*; lower triangle: *k_1_*. **B**, upper triangle: *ψ* lower triangle: *Ang_0_*. **C**, upper triangle: k_0_lk_1_; lower triangle: RTV for bevacizumab dose of 10 mg/kg. The color and asterisks indicate log_10_(p-value): ***, (p-value ≤ 0.001); **, (0.001 < p-value ≤ 0.01); *, (0.01 < p-value < 0.05).

### Determination of relationship between tumor growth parameters and response to treatment

We applied PLSR, a multivariate regression analysis, to further quantify the importance of specific tumor growth characteristics in predicting the response to anti-VEGF treatment. We used the values of *k_0_, k_1_, ψ, Ang_0_*, and *k_0_lk_1_* as inputs (predictors) and the RTV at the two dosage levels for bevacizumab (2 and 10 mg/kg) as the responses. We determined the optimal PLSR model by varying the number of components from one to six and calculating the R^2^X, R^2^Y, and Q^2^Y values (see Methods section). We also varied the number of inputs, using different combinations of the estimated parameters. The final PLSR model (i.e., the model that best predicted the responses without over-fitting) had two components and included four inputs (k_0_, *k_1_, ψ*, and Ang_0_). This PLSR model is able to accurately predict the RTV at both dosage levels (Fig 6A) and also performs well with leave-one-out cross validation (Q^2^Y = 0.92). We analyzed the PLSR model to obtain insight regarding how the four inputs relate to the outputs. The variable importance of projection (VIP) scores for the four model inputs indicate that the value of ψ is the largest contributor to predicting the RTV (Fig 6B). This suggests that the value of ψ could be used to distinguish tumors that will respond to therapy or not.

**Fig 6.**
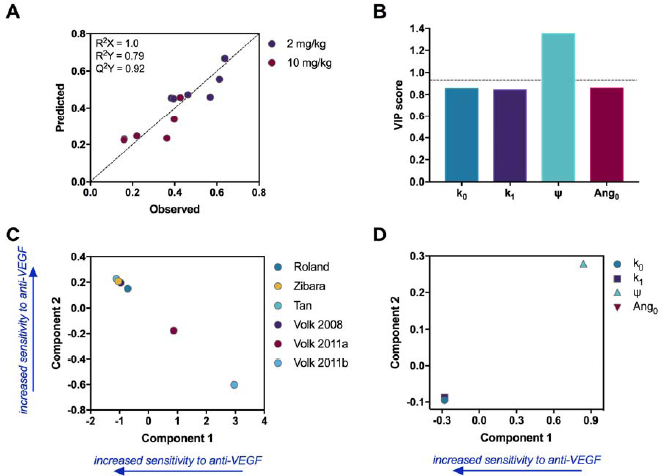
Results from multivariate analysis. PLSR analysis quantifies how the tumor growth parameters influence the response to treatment (RTV). **A**, PLSR model to predict RTV for two dosage levels of the anti-VEGF. The optimal PLSR model includes two components. Decreasing in component 1 or increasing in component 2 corresponds to higher efficacy of the anti-VEGF treatment. **B**, VIP scores for the model inputs; a score greater than one indicate variables that are important for predicting the RTV. **C**, Scores of the model output, revealing how tumors from each dataset compare in their responsiveness to treatment. **D**, Loadings of the model inputs, indicating how the model inputs (fitted parameters) correspond to sensitivity to anti-VEGF treatment.

Although the PLSR components do not explicitly correspond to a physiological variable, plotting the loadings for the inputs and outputs provides some insight into the meaning of each component. A plot of the loadings for the outputs (Fig 6C) reveals that both components capture the treatment efficacy. Here, we consider both components, as together, they account for 80% of the variance in the output. Decreasing in component 1 and increasing in component 2 corresponds to increased efficacy of the anti-VEGF treatment. Here, one of the datasets from the 2011 paper by Volk *et al*. [38], in which anti-VEG F treatment is the least effective in reducing tumor growth compared to the other datasets, has the highest loading in component 1 and lowest loading in component 2 (i.e., it appears in the lower right portion of the plot). In comparison, measurements from tumors in which anti-VEGF treatment leads to more growth inhibition appear in the upper left quadrant of the plot.

A plot of the loadings for the inputs reveals how the estimated tumor growth parameters are associated with treatment efficacy. Here, we focus only on the loadings for component 1, as this component accounts for 94% of the variance in the inputs. We find that ψ is positively correlated with low treatment efficacy, as it has a positive loading in component 1 (Fig 6D). This result suggests that a high value of *ψ* is associated with low treatment efficacy. In summary, the multivariate analysis provides a regression model that accurately predicts the relative tumor volume following anti-VEGF treatment, given the tumor growth parameters. Additionally, the analysis confirms the importance of *ψ* as a key predictor of the tumor’s response to anti-VEGF treatment.

### Effect of tumor receptor number on the response to treatment

After validating the model and investigating relationships between kinetic parameters describing tumor growth and response to treatment, we sought to investigate the effects of the tumor microenvironment. In particular, we examined the effect of neuropilin and VEGF receptor levels on relative tumor volume. VEGF receptor levels were varied from 0 to 10,000 receptors/cell, and NRP levels were varied from 0 to 100,000 receptors/cell. Using a representative set of parameters from the best fits for each dataset, we used the model to determine *T_start_* for each combination of receptor levels. We then ran the model to simulate the tumor growth for six weeks past *T_start_* to obtain the baseline control volumes. Treatment volumes were obtained by simulating twice-weekly bevacizumab injections at a dose of 10 mg/kg for six weeks after *T_start_*. The RTV values were calculated for each combination of the tumor receptor densities. The model predicts that higher neuropilin levels led to decreased treatment efficacy, especially for high VEGFR levels (Fig 7). Regardless of neuropilin density, very high treatment efficacy (low RTV) was observed when tumor cells expressed few VEGFRs. The predicted RTV values obtained using the estimated parameters from certain datasets show that neuropilin expression has a noticeable impact on the response to treatment (Fig 7A-B). In comparison, neuropilin levels seem to have a diminished impact for the Volk dataset, indicated by surfaces that are almost identical, even with drastic changes in neuropilin receptor levels (Fig 7C). In summary, the model can be used to study tumor microenvironments that provide favorable conditions for anti-angiogenic treatment. Higher receptor expression is predicted to reduce anti-VEGF efficacy, although the relationship was not uniform across all datasets, indicating the importance of accounting for specific tumor microenvironmental conditions.

**Fig 7.**
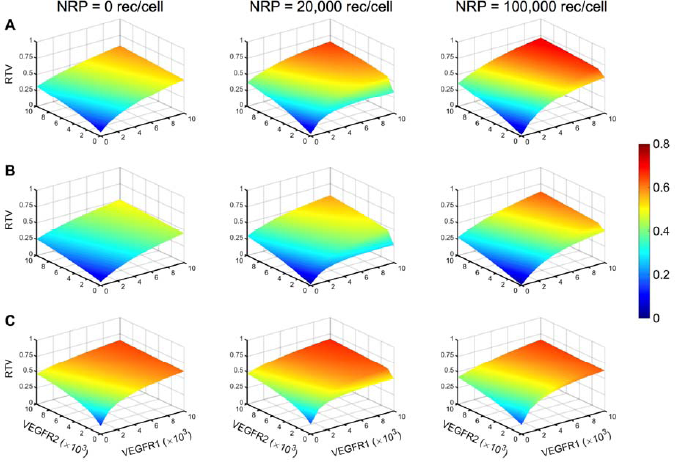
Effect of VEGF receptor expression on tumor cells. Relative tumor volume (RTV) predicted by the model using optimized parameter values obtained from fitting: **A**, Roland [34]. **B**, Zibara [35]. **C**, Volk 2008 [37], for different VEGF receptor levels on tumor cells. Neuropilin density varies: 0 receptors/cell (*left*), 20,000 receptors/cell (*center*), and 100,000 receptors/cell (*right*). Surface plots reveal the relationship between RTV and VEGFR1, VEGFR2, and neuropilin receptor density on tumor cells. The colorbar indicates the RTV value, with the same range for all panels. Red color indicates higher RTV, representing tumor conditions that are less favorable for anti-VEGF treatment.

## Discussion

We have developed a compartmental model representing tumor-bearing mice in which the tumor volume is responsive to changes in VEGF concentration. The tumor volume explicitly depends on the “angiogenic signal”, which is the signal produced when VEGF binds to its receptors on tumor endothelial cells. In this way, the model can be applied to analyze the effect of anti-VEGF treatment on xenograft tumor growth, aiding in the analysis of pre-clinical data. Kinetic parameters are obtained by fitting the model to experimental data of breast xenograft tumor growth in mice and are validated with treatment data. By including a dynamic tumor volume that explicitly depends on the concentration of VEGF-bound receptors, we address a primary limitation of our previous work.

The change in tumor volume estimated by our computational model matches experimental data used to train the model. We determine the tumor growth kinetic parameters by fitting the computational model to tumor growth measurements obtained from *in vivo* tumor xenografts. Using the fitted model, we predict the tumor volume in response to anti-angiogenic treatment targeting VEGF. These model predictions compare well with measurements of treatment response observed in six different pre-clinical *in vivo* experiments. Our approach of training the model using control data and using the optimized model to predict treatment data is a significant advantage over previous modeling work. For example, in model fitting performed by other groups, tumor growth parameters were estimated by simultaneously fitting both control and treatment groups [33] or adopted parameter values from previous models [39]. In contrast, our computational model is able to accurately predict response to anti-VEGF treatment, data not used in the fitting. This is a significant feature of our model - it is trained using control data and can reproduce the response to anti-VEGF treatment simply by introducing the drug into the blood compartment, mimicking pre-clinical mouse studies.

Using four parameters that characterize the tumor growth kinetics, our model is able to predict both untreated and treated tumor growth profiles. We note that the tumor growth equation used in previously published work [8,33] includes coefficients that characterize the killing effect of cancer drugs, including anti-angiogenic agents, on tumor growth. Since we only train the model using control data (untreated tumors), we do not include those parameters in the tumor growth equation used here. Therefore, our final model includes four parameters for the tumor growth kinetics. We did attempt to fit the data using a coefficient to describe how dependent tumor growth is on the angiogenic signal (*C_Ang_*); however, including that additional fitted parameter does not improve the fit, and the estimated value of coefficient is highly variable.

The model provides unique insight into how certain kinetic parameters that characterize tumor growth correlate with response to anti-angiogenic treatment. Our results demonstrate how the parameters describing tumor growth could be used as a predictive biomarker for treatment response. In comparison, other studies have used volume-based growth tumor kinetics as a prognostic biomarker. Lee and coworkers found that the time to progression (defined as the time it takes the tumor to grow from its nadir in volume after treatment to a progressive disease state) was significantly correlated with overall survival [11]. In other work, researchers used tumor growth kinetics to determine the efficacy of anti-angiogenic treatment [12-15]. Excitingly, our approach is highly predictive, where volumetric measurements performed prior to treatment can give insight into how the tumor might respond to an anti-VEGF agent such as bevacizumab.

We performed various analyses to quantify how the tumor growth kinetic parameters influence the response to treatment. The PLSR and statistical analyses reveal that higher *ψ* values are related to decreased treatment efficacy. The parameter *ψ* represents the transition from exponential to linear growth [8]. Thus, according to our results, anti-angiogenic treatment targeting VEGF would be more effective in tumors that have a smoother transition from exponential to linear growth (low *ψ*). The fitted parameters related to the shape of the tumor growth curve (*k_0_* and *k_1_*) were not shown to influence the response to treatment as significantly as *ψ* However, for datasets with significantly different responses to anti-VEGF treatment, the *k_0_lk_1_* ratio is also significantly different. Our statistical analyses indicate an inverse relationship between the ratio *k_0_lk_1_* and effectiveness of treatment. Simeoni *et al*. posit that *k_0_* and *k_1_* may be indicative of the initial aggressiveness of the cell line and of the response of the animal to tumor progression (i.e., immunological or anti-angiogenic response), respectively [33]. According to this interpretation, treatment would be least effective for tumors with aggressive initial growth (high *k_0_*) combined with a strong response from the animal (low k_1_). Additionally, we find that the basal angiogenic signal, *Ang_0_*, is not predictive of anti-angiogenic treatment response. This agrees with experimental results indicating that the ability of basal levels of circulating angiogenic factors to predict treatment efficacy is limited [7].

We used the model to investigate how the number of VEGF receptor and co-receptors on tumor cells influences the response to treatment. Currently, modified expression of VEGF receptors (VEGFR1, VEGFR2, or NRP1) appears to be among the most promising markers for bevacizumab treatment, though this has not consistently been replicated across different studies involving various cancer types [5]. In particular, low levels of soluble VEGFR1 expression in plasma and NRP1 expression on tumor cells are characteristics of a bevacizumab-responsive tumor [5]. Therefore, we wanted to use our model to predict the influence of tumor microenvironment on treatment efficacy. The model predicts that low levels of NRP1 or VEGFR lead to increased treatment efficacy for all datasets. The treatment is predicted to be most effective when both NRP and VEGFR have low expression; however, the effect of VEGFR levels appears to be more pronounced. That is, the treatment was still effective for high NRP levels, as long as VEGFR levels were low. These results are in agreement with other biomarker studies [9,40]. Although there was a consistent inverse relationship between receptor levels and treatment efficacy, the extent to which receptor numbers influenced the predicted relative tumor volume was not identical for all tumors. Datasets for tumors with higher *k_0_lk_1_* ratios and ψ values had higher RTV (i.e., the treatment was less effective), even for a wide range of receptor expression levels. This may indicate that intrinsic characteristics of the tumor related to its growth kinetics make anti-angiogenic treatment less effective, regardless of microenvironmental tumor conditions. As a result, solely using receptor expression as a predictive biomarker could lead to inconsistent results across tumor types.

Our model accurately predicts tumor growth profiles and matches experimental measurements. The focus of our model is on the molecular level interactions occurring between VEGF and its receptors. In our model, the number of VEGF-receptor (pro-angiogenic) signaling complexes formed directly influences tumor growth. We acknowledge that this representation of tumor growth omits the intracellular signaling pathways and corresponding cellular-level responses (i.e., proliferation and migration) involved in new blood vessel formation. However, the model does indeed capture the dynamics of tumor growth, providing a mechanistic understanding of the growth kinetics that contribute to the response to anti-VEGF treatment.

We acknowledge some assumptions and limitations that may be addressed as more quantitative data become available. We do not account for changes in tumor vascularity relative to the tumor volume. The tumor volume consists of interstitial space, vascular volume, and tumor cells. We account for tumor growth by assuming the tumor cell volume fraction increases, while the interstitial space volume fraction decreases, and the relative proportion of the vascular volume is constant (see Methods section for more detail). This means that the tumor vascularity does change as the overall tumor volume grows, but it remains in the same proportion relative to the whole tumor volume. Furthermore, we do not simulate remodeling of the blood compartment or changes in vascular permeability in response to anti-VEGF treatment.

However, experimental data show a decrease in microvessel density following bevacizumab treatment [41], and incorporating this observation would enhance the model. Additionally, anti-angiogenic treatment promotes normalization of the vasculature, which allows for more efficient delivery of chemotherapy to the tumor [42]. Accounting for changes in the microvascular density would allow us to simulate combination treatments that include chemotherapy and anti-angiogenic agents. Unfortunately, there is a lack of robust time-series data that can be used to predict changes in vascular density with treatment. This limitation may be addressed as additional quantitative measurements are published.

The model is highly successful in capturing the growth kinetics of exponential or linear growth curves. However, the model does not accurately predict sigmoidal tumor growth. The equation governing tumor growth used in our model is based on the foundational work of Simeoni *et al.*, who adapted a Gompertz model of tumor growth to investigate both the exponential and linear phases of growth [33]. Although this makes the tumor growth equation more flexible, it also limits the ability to simulate an eventual plateau in growth. The model’s inability to capture sigmoidal growth was particularly apparent when fitting the Volk datasets [37,38]. However, we have focused on exponential growth, as it has been implemented in many other mathematical models [43,44] and shown to accurately fit tumor growth data [45]. Expansion of the tumor growth equation can be added in future studies.

### Concluding thoughts

We constructed a computational model that simulates the kinetics of VEGF binding to its receptors and the influence of VEGF-bound receptor complexes on tumor volume in tumor-bearing mice. This model is a useful tool in the analysis of pre-clinical data. The model matches control tumor growth data used for fitting, and it was validated using data not used in fitting. The validated model accurately predicts the tumor growth upon administration of anti-angiogenic treatment that targets VEGF. Recently published imaging studies indicate that tumor growth kinetics are prognostic biomarkers for treatment efficacy. Interestingly, the fitted parameter values estimated in the present study point to the possibility of using tumor growth kinetics as a predictive biomarker for anti-angiogenic treatment. This model also helps to elucidate why biomarker candidates such as expression of VEGF receptors on tumor cells may not be reliable for all tumors. Although the model predicts that receptor levels influence response to treatment, the effects are not uniform across all of the experimental datasets we analyzed. Thus, our modeling work lays the foundation for future studies to investigate the importance of tumor growth kinetics as a predictive and specific biomarker and can accelerate the discovery of biomarker candidates in pre-clinical studies.

## Materials and Methods

### Computational Modeling

#### Compartmental model

In this work, we expand on our previous three-compartment model [27] by including VEGF-mediated tumor growth. We briefly describe the full model and detail the new additions that are the focus of this work. The model is comprised of three compartments representing the whole mouse: normal tissue (assumed to be skeletal muscle), blood, and tumor (Fig 8). The model includes human and mouse VEGF isoforms: human isoforms (VEGF_121_ and VEGF_165_) are secreted by tumor cells, and mouse isoforms (VEGF_120_ and VEGF_164_) are secreted by endothelial cells in the normal, blood, and tumor compartments and muscle fibers in the normal tissue. Geometric characteristics, receptor densities, kinetic parameters, and transport rates are all detailed in our previous paper [27].

**Fig 8.**
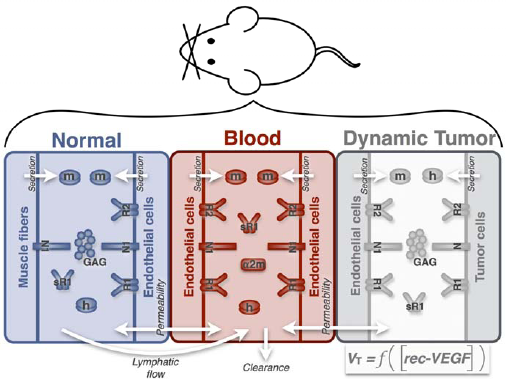
Model schematic. The computational model includes three compartments: normal tissue, blood, and tumor volume. The compartments are connected via lymphatic flow from the interstitial space in the normal tissue to the blood and transendothelial macromolecular permeability. Molecular species include human and mouse VEGF isoforms, VEGF receptors and co-receptors (including the soluble receptor VEGFR 1, sR1), and the protease inhibitor α-2-macroglobulin (a2m). Glycosaminoglycan (GAG) chains represent the extracellular matrix. The volume of the tumor depends on the concentration of receptor-bound VEGF complexes on tumor endothelial cells (denoted as [rec-VEGF]).

#### Tumor volume and growth

Previously, we assumed the tumor volume increased exponentially with time, based on measurements from tumor xenografts [27]. Under that assumption, cancer treatment, including anti-angiogenic therapy, has no effect on tumor growth. In the present study, we address that limitation by introducing an equation for tumor growth wherein the volume of the tumor compartment is dependent on the “angiogenic signal” (*Ang*) produced when VEGF binds to its receptors on endothelial cells in the tumor.

The tumor compartment is assumed to consist of cancer cells, endothelial cells (vascular volume) and interstitial space, each of which has a defined volume fraction (i.e., volume relative to the total tumor volume). Our previous model assumed that as the total tumor volume increased, the relative proportions of cancer cells, vascular space, and interstitial space remain constant. Here, we still have the volume fraction for the vascular space remaining constant, based on a range of experimental data [46-48]. However, we used results from a recent imaging study to account for an increase in the relative volume of cancer cells as the tumor volume increases. Christensen and coworkers measure how tumor cell density increases as the tumor grows by tracking cancer cells in xenograft tumors in rats using near near-infrared (NIR) fluorescence dyes [49]. The authors quantify the fluorescence intensity in a tumor and use it to estimate the number of cancer cells as the tumor grows over time. The estimated cell count was normalized by the tumor volume to obtain the number of cancer cells per unit volume of tumor tissue as the tumor grows. We extracted the values obtained by Christensen and coworkers for MDA-MB-231 tumors and converted them to the cancer cell volume fraction using the volume of tumor cells, as we have done in our previous work [27]. Therefore, we have been able to incorporate into our model an increase in the cancer cell volume fraction over time. Assuming a tumor cell volume of 905 μm^3^, based on our previous examination of the literature [27], we developed expressions describing the decay of interstitial space during tumor growth. We found that the relative decrease in interstitial space during tumor growth was adequately modeled by exponential decay. The equations for how the relative volume of the interstitial space varies with total tumor volume are given in Supplemental File S5.

### Data Extraction

Data from six independent *in vivo* published experimental studies of MDA-MB-231 xenograft tumors in mice were used for parameter fitting and validation [34-38]. The six datasets included growth profiles for untreated tumors (control), as well as tumors treated with the anti-VEGF agent bevacizumab. Experimental data was extracted using the WebPlotDigitizer program (http://arohatgi.info/WebPlotDigitizer). The numerical values are provided in Supplemental File S3.

### Parameter estimation

#### Model training

We first fit *k_0_*, k_1_ *ψ* and *Ang_0_*(“free parameters”) using the control tumor growth profiles for each dataset. Fitting was performed using the *lsqnonlin* function in MATLAB to minimize the sum of squared residuals (SSR):

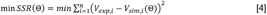

where *V_exp,i_* is the ith experimentally measured tumor volume, *V_sim,i_* is the ith simulated volume at the corresponding time point, and *n* is the total number of experimental measurements. The minimization is subject to Θ, the set of upper and lower bounds on each of the free parameters. We found that weighting the residual by the experimental measurement biased the error towards early data points and reduced the model’s ability to fit the full course of tumor growth. Therefore, we minimized the residual, with no weighting, to fit the model to the experimental data.

We performed the parameter fitting 20 times for each dataset. To attempt to arrive at a global minimum for the error, we initialized each fitting run by randomly selecting a value for the free parameters within the specified upper and lower bounds. The bounds were set such that the range for each parameter was at least one order of magnitude: 10^-8^ to 10^-2^ for *k_0_* and *k_1_* 0.1 to 50 for *ψ* and 10^-16^ to 10^-14^ for *Ang_0_*. After performing the model fitting, we used the SSR to identify the optimal parameters. Parameter sets with the smallest errors were taken to be the “best” fits and were used for subsequent statistical analysis. The number of “best” parameter sets varied between datasets and ranged from 12 to 16 parameter sets. We first tested to see whether there were significant effects of the experimental data being fit on the estimated parameters values using one-way non-parametric ANOVA. This method makes no assumptions about the distributions of parameter values and tests whether samples originate from a common distribution. We then performed post-hoc analyses to make pairwise comparisons using the Kruskal-Wallis test. We corrected for multiple comparisons using statistical hypothesis testing (Dunn’s test). All statistical analyses were performed using GraphPad Prism.

Two of the experimental datasets contained at least three data points prior to administration of treatment [34,38]. These points were used in a separate model fitting to see whether limiting the data used for model training to only pre-treatment measurements could generate a fitted model that still accurately predicts the response to anti-angiogenic treatment.

#### Model validation

After fitting the control data, we validated the estimated parameters with data not used in the fitting. We applied the fitted model to simulate anti-angiogenic treatment and compared the predicted tumor growth profile to the experimental measurements for the treatment cases. Here, we simulated the dosing regimens used in each experiment with the same optimized parameters obtained by fitting the control data. For each dataset, we simulated intravenous injections lasting for one minute. All six experimental studies administered bevacizumab bi-weekly; however, the dosage varied between the studies. The dosing regimens are given in Supplemental File S6. The binding affinity and clearance rate for bevacizumab were obtained from experimental studies in which VEGF was immobilized on a flow cell (FC) and bevacizumab was injected over the FC at varying concentrations [50]. Based on those measurements, the binding affinity was set to 4456 pM (*k_on_* = 5.4 × 10^4^ M^-1^ s^-1^; *k_off_* = 2.19 × 10^-5^ s^-1^), and 5.73 × 10^-7^ s^-1^ was used for the anti-VEGF clearance rate.

### Partial least squares regression analysis

Partial least squares regression (PLSR) modeling was used to determine the relationship between the fitted parameters characterizing tumor growth kinetics (inputs) and the response to treatment given by the RTV value (output). PLSR modeling seeks to maximize the correlation between the inputs and outputs. To accomplish this, the inputs and outputs are recast onto new dimensions called principal components (PCs), which are linear combinations of the inputs. The regression coefficients estimated by PLSR describe the relative importance of each input. Quantitative measures from the PLSR modeling, including the loadings and scores, provide some insight into the biological meaning of the PCs [51]. Additionally, we use the estimated regression coefficients to determine each input’s contribution across all responses. This metric is given by the “variable importance of projection” (VIP) for each predictor. The VIP value is the weighted sum of each input’s contribution to all of the responses. As such, the VIP values indicate the overall importance of the predictors. VIP values greater than one indicate variables that are important for predicting the output response.

For our analysis, the input matrix was 6 rows × 4 columns, where the 6 rows correspond to the best fit for each of the six datasets, and the 4 columns consisted of the estimated free parameters (k_0_, *k_1_, ψ*, and *Ang_0_*). The output matrix was 6 rows × 2 columns, where the rows corresponds to the predicted RTV using the best fit for each of the six datasets, and the columns are the two treatment doses (2 and 10 mg/kg). We used two metrics to evaluate the model fitness: R^2^Y and Q^2^Y, which each have a maximum value of 1. The R^2^Y value indicates how well the model fits the output data. The Q^2^Y metric specifies how much of the variation in the output data the model predicts [52], and values greater than 0.5 indicate that the model can predict data not used in the fitting. We performed PLSR modeling using the nonlinear iterative partial least squares (NIPALS) algorithm [53], implemented in MATLAB (Mathworks, Inc.).

### Numerical implementation

All model equations were implemented in MATLAB using the SimBiology toolbox. The final model is provided as SBML (Supplemental File S7). Parameter fitting was performed using the *lsqnonlin* function MATLAB. GraphPad Prism was used to run statistical analyses on parameter values.

## Acknowledgements

The authors thank members of the Finley research group for critical comments and suggestions. The research was supported by NSF CAREER award 1552065 (awarded to SDF) and a USC Provost’s Undergraduate Research Fellowship (awarded to TDG).

## Author Contributions

SDF designed the research. TDG, ADA, and SDF performed the simulations. TDG and SDF analyzed the data and wrote the manuscript.

## Supporting Information

**S1 Table. Experimental data extracted from published papers.** This table lists the experimental data used for model fitting.

**S2 Figure. Comparison of error from model fitting.** We fit published experimental data for xenograft tumor volumes using two different models to characterize the volume of the tumor (equations (1) and (2), termed Tumor Growth Models 1 and 2, respectively). The sum of the squared residual (SSR) for the best fits using either equation for each of the six datasets is plotted. **A**, Roland [34]. **B**, Zibara [35]. **C**, Tan [36]. **D**, Volk [37]. **E**, Volk 2011a [38]. **F**, Volk 2011b [38]. The analysis shows that adding a fifth fitted parameter (*C_Ang_* in Tumor Growth Model 2) does not significantly lower the error, indicating that Model 2 does not provide a better fit to the experimental data. The horizontal bar indicates the median of the best fits obtained from fitting the model to each dataset.

**S3 Figure. Estimated model parameters obtained from fitting Tumor Growth Model 2.** The estimated parameter values from the best fits are plotted for each dataset when using Tumor Growth Model 2, equation (2), to calculate the tumor volume. This model includes an additional fitted parameter (*C_Ang_*), compared to Tumor Growth Model 1. **A**, *k_0_*. **B**, *k_1_*. **C**, *ψ* **D**, *Ang_0_*. **E**, *k_0_lk_1_*. **F**, *C_Ang_*. The horizontal bar indicates the median of the best fits obtained from fitting the model to each dataset.

**S4 Figure. Predicted response to anti-VEGF treatment.** The model was used to simulate bevacizumab treatment at a dose of 2 mg/kg. The relative tumor volume (RTV) predicted by the model is shown. Horizontal bar indicates the median of the predicted RTV for the best fits from each dataset.

**S5 Table. Equations describing change in relative volume of the interstitial space.** This table presents the equations for how the relative volume of the interstitial space changes as a function total tumor volume. This equation is unique for each of the datasets investigated.

**S6 Table. Experimental treatment conditions.** This table describes the experimental conditions for anti-VEGF treatment, taken from published studies used in model fitting.

**S7 Dataset. Model file.** This file contains the computational model in SBML format.

